# Virtual EEG-electrodes: Convolutional neural networks as a method for upsampling or restoring channels

**DOI:** 10.1101/2020.04.20.049916

**Authors:** Mats Svantesson, Håkan Olausson, Anders Eklund, Magnus Thordstein

**Affiliations:** Department of Clinical Neurophysiology, University Hospital of Linköping, Sweden; Center for Social and affective Neuroscience, Linköping University, Sweden; Center for Medical Image Science and Visualization, Linköping University, Sweden; Department of Biomedical Engineering, Linköping University, Sweden; Division of Statistics & Machine Learning, Department of Computer and Information Science, Linköping University, Sweden; Department of Biomedical and Clinical Sciences, Linköping University, Sweden

**Keywords:** Deep learning, Convolutional neural networks, Electroencephalography, Signal reconstruction, Spatial upsampling, Spherical spline interpolation

## Abstract

**Background:** In clinical practice, EEGs are assessed visually. For practical reasons, recordings often need to be performed with a reduced number of electrodes and artifacts make assessment difficult. To circumvent these obstacles, different interpolation techniques can be utilized. These techniques usually perform better for higher electrode densities and values interpolated at areas far from electrodes can be unreliable. Using a method that learns the statistical distribution of the cortical electrical fields and predicts values may yield better results.

**New Method:** Generative networks based on convolutional layers were trained to upsample from 4 or 14 channels or to dynamically restore single missing channels to recreate 21 channel EEGs. 5,144 hours of data from 1,385 subjects of the Temple University Hospital EEG database were used for training and evaluating the networks.

**Comparison with Existing Method:** The results were compared to spherical spline interpolation. Several statistical measures were used as well as a visual evaluation by board certified clinical neurophysiologists. Overall, the generative networks performed significantly better. There was no difference between real and network generated data in the number of examples assessed as artificial by experienced EEG interpreters whereas for data generated by interpolation, the number was significantly higher. In addition, network performance improved with increasing number of included subjects, with the greatest effect seen in the range 5 – 100 subjects.

**Conclusions:** Using neural networks to restore or upsample EEG signals is a viable alternative to interpolation methods.

## 1 Introduction

In clinical routine, EEG is assessed by visual inspection by a trained human interpreter. Ongoing work strives to develop methods for automating the analysis of EEG (Acharya et al., 2013; Roy et al., 2019). However, it is likely that the analysis will continue to be visual for the foreseeable future and that there will be a long period of transition, where visual and automated analysis will be performed in parallel, before the analysis becomes fully automated. Developing methods to enhance the visual assessment is thus motivated.

EEG is used increasingly for long term monitoring of cerebral function, e.g., to detect seizures or status epilepticus (Kubota et al., 2018). The number of scalp electrodes used in this setting is often reduced compared to a standard EEG (e.g., 4 instead of 21 electrodes), due to limitations in resources and practical problems with maintaining signal quality for longer time periods (Hera et al., 2017). A low electrode density gives a low spatial resolution, making the recordings harder to assess compared to standard EEGs since identification and classification of wave phenomenon often depend on spatial patterns.

Common methods used for scalp potential interpolation are nearest-neighbor and splines (Fletcher et al., 1996). For instance, the commercial EEG analysis software Curry (Compumedics Neuroscan, Dresden, Germany) has a nearest-neighbor implementation, the Python library MNE (Gramfort et al., 2013) and Matlab toolbox EEGLAB (Delorme and Makeig, 2004) uses spherical splines as default method, and FieldTrip toolbox (Oostenveld et al., 2011) has weighted average of nearest neighbors as default with the option of spherical splines. Spline techniques seem to perform better than nearest-neighbor with a tendency for spherical splines to perform best (Perrin et al., 1987; Perrin et al., 1989; Soong et al., 1993). Bilinear and bicubic interpolation are other common methods used for interpolation (Koles and Paranjape, 1988). There seem to be few studies were systematic comparisons are made with splines methods, but thin plate splines have been shown to perform better than bilinear interpolation in topographic mapping (Satherley et al., 1996). It has been concluded that ‘…adequate electrode density is more important than the method used’ and that there is a risk of large interpolation errors in areas distant to electrodes (Fletcher et al., 1996). It has also been claimed that the electrode density used clinically, i.e., the international 10-20 system (Jasper, 1958), is too low for interpolation (Soong et al., 1993).

When deep learning applications are used for EEG analysis, convolutional neural networks (CNNs) are most common, used in 40 % of the papers (Roy et al., 2019). Some of these studies investigate the generation of synthetic EEG data, with the majority of networks implemented as generative adversarial networks (GANs). Most of the studies concern the generation of synthetic signals for data augmentation and have been shown to provide realistic signals and improve classification of data (Pascual et al., 2019; Aznan et al., 2019; Zhang and Liu, 2018; Hartmann et al., 2018; Luo and Lu, 2018; Qiu and Zhao, 2018). Wasserstein GANs (WGANs) have been used for temporal upsampling, also demonstrating an improvement in classification tasks (Luo et al., 2020). Only a few studies are related directly to spatial upsampling, relevant to the work here presented. WGANs have been used to upsample from 8 or 16 to 32 channels (Corley and Huang, 2018). This result was achieved using the training dataset ‘V’ of Berlin Brain Computer Interface Competition III (Millan, 2004) consisting of 2,141 s of 32 channel EEG data from 3 subjects. The method produced a reduction of the mean absolute error by one to two orders of magnitude compared to bicubic interpolation. CNNs have been used to recreate 64-channel EEG from as low as 4 channels (Kwon et al., 2019). However, the inputs to the networks were first linearly interpolated to 64 channels. Training was performed with simulated EEG data (640 examples for training) and experimental EEG data from an auditory task (596 examples from 5 subjects for training). The results were compared to the interpolation used as input, showing a lower mean square error and higher correlation with the original data; in addition, source localization using recreated signals was fairly adequate whereas it failed with interpolated data.

It is usually necessary to train deep neural networks with large amounts of data (Sun et al., 2017). When training for EEG processing, utilizing data from an adequate number of individuals may be important for generalization but this has not been thoroughly investigated. Using 1,500 ms examples of 128-channel EEG data from up to 30 subjects in a cognitive task (the number of examples was not reported), an improvement in classification accuracy was seen with increasing number of subjects, most notably when more than 15 subjects were included (Völker et al., 2018).

In this study we further investigate the use of CNNs to upsample the electrode density or to restore signals of EEG and compare the results to a standard method used in common software packages for EEG processing, spherical splines. We evaluate the signal quality visually as well as with several objective measures to assess the closeness to real signals. Most previous studies have used a limited amount of data and few subjects, but here we use a large data set of real EEG and also test to what extent training the networks with different number of subjects affects the results.

## 2 Materials and methods

### 2.1 Software and hardware

The computers used the operative system Ubuntu version 18.04.2 LTS. The networks were developed in Python (version 3.6.5) using the API Keras (version 2.2.4) and programming module TensorFlow (version 1.10.1). Random seed was set to 12,345 for the libraries ‘random’ and ‘numpy.random’.

The Python library ‘pyEDFlib’ (Nahrstaedt and Lee-Messer, 2017) was used to extract EEG data. Extracting subject information, i.e., age and gender, was not possible with the ‘pyEDFlib’, so MATLAB R2018a was used with a downloaded script (Shapkin, 2012). The MNE-Python library (Gramfort et al., 2013) was used to perform spherical spline interpolation.

Two computers were used for the study, equipped with: 1) Intel Xeon E5 1620V4 Quad Core 3.5 GHz processor, 96 GB 2,400 MHz RAM and Nvidia Quadro P5000 GPU; 2) Intel Core i7-6800K Hexa Core 3.4 GHz, 96 GB 2,133 MHz RAM, Nvidia Quadro P6000 and Nvidia Titan X.

### 2.2 EEG data

The EEG data used for this study was acquired from the published data base created at the Temple University Hospital (TUH), Philadelphia (Obeid and Picone, 2016). The TUH EEG Corpus (v1.1.0) with average reference was used (downloaded during 17-21 January 2019).

The recording electrodes were positioned according to the international 10-20 system (Jasper, 1958). A full EEG montage here is defined as consisting of 21 electrodes Fp1, F7, T3, T5, Fp2, F8, T4, T6, F3, C3, P3, O1, F4, C4, P4, O2, A1, A2, Fz, Cz and Pz; here given in the order used for visual analysis. Many recordings had more electrodes but only those listed here were used. The data was not re-referenced.

A total of 1,385 (♂/♀: 751/634) subjects with 11,163 (♂/♀: 5,648/5,515) recordings, corresponding to 5,144 hours, was extracted from the data set. Ages varied from 18 to 95 years with a mean age of 51±18 years. The data contained normal and pathological as well as wake and sleep recordings of unknown distributions. All original recordings were in European Data Format (EDF), unfiltered and sampled at 256 Hz. An arbitrary lower limit of recording length of 300 seconds was used to ensure that each recording would offer some variance during training.

#### 2.2.1 Data pre-processing

All data was high pass filtered at 0.3 Hz and low pass filtered at 40 Hz using second-degree Butterworth filters; the bandwidth was chosen according to our clinical preference. A 60 Hz notch filter was used to remove residual AC-noise. Filtering was applied with zero phase shift.

The data were divided into a training, validation and test set (80, 10 and 10 percent distribution, respectively). To ensure that the sets were disjoint with regard to subject, each subject was randomly assigned to one of the sets according to the distribution, resulting in 1,114, 129 and 142 subjects for each data set.

The training data consisted of 1,114 subjects providing, in total, 3,976 hours of data. To evaluate the effect of the number of subjects used, networks were also trained with subsets of 5, 15, 30, 50, 100 and 500 randomly chosen subjects. This resulted in subsets consisting of 11, 37, 152, 159, 414 and 1,694 hours of data, respectively.

Data examples with a duration of 10 s were generated randomly during training, validation and testing, see section 2.3.3; to avoid very large artifacts, drawn examples with maximum absolute amplitude over 500 μV were discarded. The first and last 40 s of each recording was also discarded since these parts sometimes had more artifacts or did not contain any cortical signal (usually a low frequency square wave). This time interval was arbitrarily chosen from visual assessment of recordings, did not exclude all epochs without cortical signal and also resulted in discarding epochs with good data. No other measures were taken to reduce artifacts.

Various approaches to normalization were evaluated; none, normalization to an amplitude- or a standard deviation of one. The latter option performed best, most notably a much better estimation of amplitudes. Since the data were high pass filtered, the mean was close to zero on average.

#### 2.2.2 Ethical considerations

The HIPAA Privacy Rule was followed when the TUH EEG Corpus was created (Obeid and Picone, 2016). Since the used data was from a published data base with no subject information and therefore no way of tracing the data back to the subjects, no further approval from an ethic committee was deemed necessary for this study.

### 2.3 The networks

#### 2.3.1 Architecture

In this study, three generative networks were trained for processing 10 s examples of EEG data in slightly different ways. The first (GN1) upsampled from 4 to 21 electrodes, the second (GN2) upsampled from 14 to 21 electrodes and the third (GN3) could recreate any of the 21 electrodes. The networks shared the same general architecture, the only essential difference was in the number of input and output channels. The electrodes used as inputs were not recreated meaning that GN1 had 4 inputs and 17 outputs, GN2 had 14 inputs and 7 outputs and GN3 had 21 inputs and 1 output (see Table 1). The function of the GN3 network differed since it was trained to dynamically replace anyone of the electrodes, meaning that it also had to learn to detect which channel it was supposed to recreate.

**Table 1.** Input and output electrodes of the three networks.

| Network | Input electrodes | Output electrodes |
| --- | --- | --- |
| GN1 | F3, P3, F4, P4 | Fp1, F7, T3, T5, Fp2, F8, T4, T6,<br>C3, O1, C4, O2, A1, A2, Fz, Cz, Pz |
| GN2 | Fp1, F7, T5, Fp2, F8, T6, C3, O1, C4, O2,<br>A1, A2, Fz, Pz | T3, T4, F3, P3, F4, P4, Cz |
| GN3 | Fp1, F7, T3, T5, Fp2, F8, T4, T6, F3, C3,<br>P3, O1, F4, C4, P4, O2, A1, A2, Fz, Cz, Pz* | One randomly chosen of the inputs |

The given electrode order is the same as the order they are represented in the data. \*) The randomly chosen electrode is blocked as input and replaced with low amplitude noise.

The networks were based on convolutional and deconvolutional layers, and except for the last layer, all layers were followed by a leaky rectifying linear unit (Maas et al., 2013). Convolutions and deconvolutions were performed separately for temporal and spatial dimensions.

The networks had three principal parts: First part, a temporal encoder consisting of four convolutional layers, using strides of two and doubling the number of filters for each layer. Second, a spatial analysis and upsampling performed by convolution with a filter size corresponding to the number of input electrodes, and then deconvoluting to the right number of output electrodes. Third, a temporal decoder of four deconvolutional layers with strides of two and a finishing convolutional layer to merge the filters resulting in the desired size (see Table 2). With a duration of 10 s, this corresponded to 2560 samples per electrode for in- and output. GN1 and GN2 had 6,546,241 and GN3 had 6,808,385 parameters, all trainable.

**Table 2.**
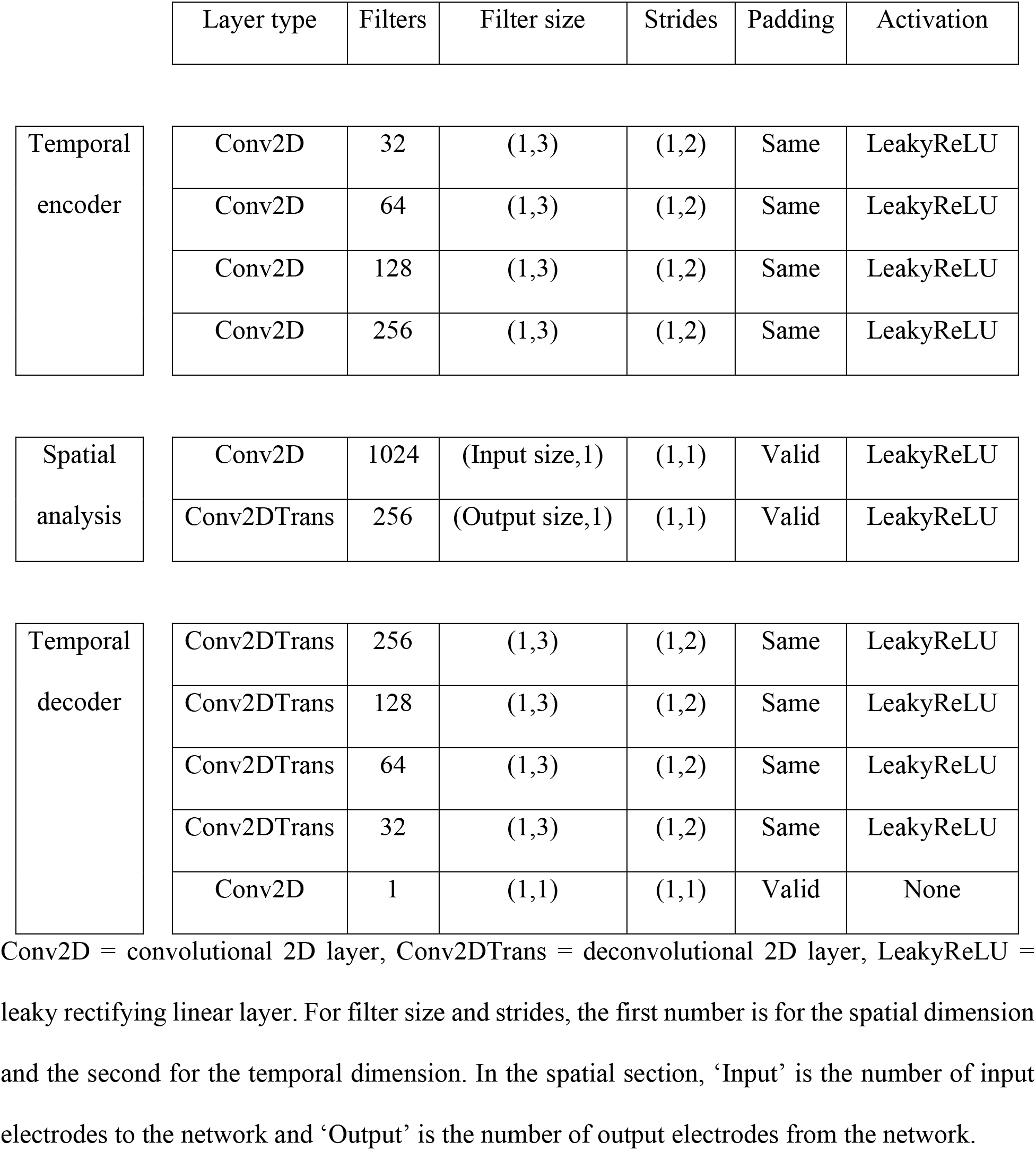
Overview of the generative network.

|  | Layer type | Filters | Filter size | Strides | Padding | Activation |
| --- | --- | --- | --- | --- | --- | --- |
| Temporal<br>encoder | Conv2D | 32 | (1,3) | (1,2) | Same | LeakyReLU |
|  | Conv2D | 64 | (1,3) | (1,2) | Same | LeakyReLU |
|  | Conv2D | 128 | (1,3) | (1,2) | Same | LeakyReLU |
|  | Conv2D | 256 | (1,3) | (1,2) | Same | LeakyReLU |
| Spatial<br>analysis | Conv2D | 1024 | (Input size,1) | (1,1) | Valid | LeakyReLU |
|  | Conv2DTrans | 256 | (Output size,1) | (1,1) | Valid | LeakyReLU |
| Temporal<br>decoder | Conv2DTrans | 256 | (1,3) | (1,2) | Same | LeakyReLU |
|  | Conv2DTrans | 128 | (1,3) | (1,2) | Same | LeakyReLU |
|  | Conv2DTrans | 64 | (1,3) | (1,2) | Same | LeakyReLU |
|  | Conv2DTrans | 32 | (1,3) | (1,2) | Same | LeakyReLU |
|  | Conv2D | 1 | (1,1) | (1,1) | Valid | None |

Conv2D = convolutional 2D layer, Conv2DTrans = deconvolutional 2D layer, LeakyReLU = leaky rectifying linear layer. For filter size and strides, the first number is for the spatial dimension and the second for the temporal dimension. In the spatial section, ‘Input’ is the number of input electrodes to the network and ‘Output’ is the number of output electrodes from the network.

#### 2.3.2 Hyperparameters

The choice of four layers and a stride two were empirical; it most consistently gave visually pleasing results (assessed by specialist in clinical neurophysiology) and offered shorter training times compared to larger nets. The validation losses were usually about the same for nets with two to seven convolutional layers, but more layers showed a tendency for high frequency artifacts, and fewer layers to have inaccurate phases of waveforms. Inaccurate phases of waveforms were also seen more often when using a stride of one. The formation of features was more evident for a stride of two compared to one, with the stride of one giving an impression of just passing the signal through the network.

Training with mini batches, up to a size of 128, did not improve results; this was tested with and without batch normalization. Dropout layers did not improve results. Leaky rectifying linear units performed better than rectifying linear units; an alpha of 0.2 was used.

Optimization was performed with Adam (Kingma and Ba, 2015); no other algorithm was tested. Learning rates above 10^−4^ showed an instability during training, so for the final run this rate was chosen. Using a linear decay for the learning rate did not improve results. Decay parameters β_1_ = 0.5 and β_2_ = 0.99 were used. Mean absolute error was used as the loss function. Least square error was tested as well as versions of mean absolute and least square error with an electrode gradient that gave a larger penalty for errors of electrodes with consistently higher errors relative to other electrodes. All weights of the convolutional and deconvolutional layers were initialized according to default setting, i.e., the kernel as a Glorot uniform distribution and the bias set to zero. No further regularization was used in the networks.

#### 2.3.3 Training schedule

Training were performed with overlapping examples, where an example could start at any sample point within a recording allowing extraction of a 10 s example. An epoch was defined as training corresponding to one example from all (1,114) subjects one time. For the subsets, this meant training 223, 74, 37, 22, 11 and 2 times with each subject per epoch.

The training order of the subjects was randomized for each epoch. For subjects with several EEG recordings, one was chosen at random. A start position defining a 10 s example was then randomly selected. If a drawn example from a subject was discarded due to high amplitude, attempts were made to redraw a new example with acceptable amplitude up to 100 times. If the subject had several recordings and the number of attempts were exceeded, then the next recording was tried and so on. If the number of attempts was exceeded for all recordings, no training took place for that subject during that epoch.

When training the GN3 network, 1 of the 21 input electrodes was randomly selected and the input was replaced by noise of normal distribution with mean 0 and a standard deviation of 0.1.

To decide the number of epochs, all networks were trained for up to 1,000 epochs, the epoch of minimum loss for the validation data was identified (Table 3), which was then used for the final run. For sessions with subject count below 50, a deterioration of performance was seen after some interval of training but in the other cases a stagnation of improvement was seen at some point but with no sign of overfitting.

**Table 3.** The number of epochs of training for the networks GN1, GN2 and GN3 when evaluating the effect of number of subjects in training data.

| Number of subjects | 5 | 15 | 30 | 50 | 100 | 500 | 1,114 |
| --- | --- | --- | --- | --- | --- | --- | --- |
| GN1 | 60 | 100 | 100 | 400 | 1,000 | 1,000 | 1,000 |
| GN2 | 100 | 100 | 100 | 400 | 400 | 1,000 | 1,000 |
| GN3 | 100 | 100 | 100 | 400 | 400 | 1,000 | 1,000 |

One epoch of training usually took 40 – 60 s, depending on GPU and network. Evaluation during training to monitor overfitting were done at the end of each epoch with 1,000 examples from training and validation data, respectively, which added 30 – 60 s. Total training time for 1,000 epochs with evaluation at each epoch could be 15 – 30 hours.

### 2.4 Spherical spline interpolation

As a baseline for the evaluation of the networks, spherical spline interpolation was used to recreate the same signals of the test data set as the convolutional networks. This method was chosen since it seems to be one of the best performing of the standard interpolation techniques (Perrin et al., 1987; Perrin et al., 1989; Soong et al., 1993) and is used in several common EEG processing softwares. The method consists of fitting a spherical surface to the known values of some electrodes and their corresponding spatial positions. Interpolated values can then be extracted from the resulting deformation of the surface (Freeden, 1984).

### 2.5 Evaluation

5,000 randomly chosen examples of the test data set were used for the final evaluation. The networks were evaluated in several ways using different methods and measures. The measures were: mean absolute error (MAE), correlation coefficient (R), coherence (C), cross spectral phase (P), Fréchet distance (FD) and Kullback-Liebler divergence (KL). They all reflect some aspect of the closeness or distance between generated and original data. Before calculating the measures, all data were renormalized to reflect real amplitudes (μV).

MAE was calculated straight forwardly by taking the absolute value of the difference between the original and recreated value *x_i_* and *y_i_* at each data point *i* and then taking the mean of all values *N* as 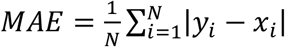.

R was calculated between the original and recreated signals with the Spearman’s rank correlation coefficient. Spearman’s method was used since the relatively high occurrence of artifacts and biological signal transients resulted in a non-Gaussian distribution.

The coherence C was calculated as 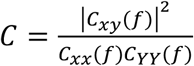, where *C_xy_*(*f*) is the Fourier transform of the cross-correlation, *C_xx_*(*f*) and *C_yy_*(*f*) are the Fourier transforms of the auto-covariances of the signals *x* and *y*. The phase P was calculated as the angle between the real and imaginary part of *C_xy_*(*f*). Values for frequencies up to 40 Hz were used.

FD is a measure of distance between two curves which also reflects differences in shape and was calculated using recursive algorithms (Eiter and Mannila, 1994). The analysis was limited to 500 examples of randomly chosen 2 s intervals due to the long processing times of the recursive algorithm.

KL is an entropy-based measure of the difference between the distributions *P* and *Q* of two data sets (Kullback and Liebler, 1951), where *P* in this case were the original data and *Q* the recreated data. It is calculated as 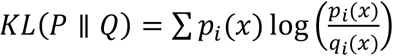. The distributions were calculated by binning the data in the interval (−500,500) μV using bins of size 5 μV.

### 2.6 Comparison with spherical spline interpolation

Signals recreated by the networks were compared to the corresponding signals recreated by spherical spline interpolation. MAE, R, C, P, FD and KL were calculated for each paired data point of original and recreated signals. Wilcoxon signed rank test was used to assess differences in the median value of the measures; for P the absolute value was used.

Speed tests were performed where a comparison of network prediction and interpolation calculation time for 5,000 examples were measured. The tests were executed on a computer with an Intel Xeon E5 1620V4 Quad Core 3.5 GHz processor, 96 GB 2,400 MHz RAM and Nvidia Quadro P5000 GPU.

### 2.7 Visual assessment of data

Blinded tests were performed where five senior consultants in clinical neurophysiology (from our clinic, not otherwise involved in this project) visually assessed randomly displayed real, network generated or interpolated EEG data. Their task was to determine whether each presented example was real, or computer generated. The purpose was to validate the visual quality, i.e., that wave morphology and fields appeared credible.

125 of the 5,000 test examples were chosen at random. The 125 examples were in turn randomized as either being represented by the original signals, the signals recreated by the GN1 network or by interpolation. The same randomization was used for all raters. A Python script was developed which displayed one example at a time. For each example, the rater had to press one of two buttons in the GUI to rate it as real or artificial. Pressing a button automatically loaded the next example with no possibility of going back. The first 25 examples were used for practice, after which it was verified that the rater had understood the task, and the remaining 100 examples were then used for the test. Each neurophysiologist had 20 minutes to complete the test part. The data was displayed in average montage, the amplitude could be altered but no other adjustments were possible.

A Chi-squared test was used to assess differences. The median rating of each example was used, and testing was done pairwise between data types.

### 2.8 The effect of the number of subjects

The effect of the number of subjects, also indirectly reflecting the amount of data, used for training were evaluated with MAE, R and C. A Kruskal-Wallis H test was used for multiple comparisons and when the null hypothesis could be rejected, post hoc analyses were performed by Dunn’s multiple comparison test using an adjusted p-value corresponding to 0.05.

## 3 Results

### 3.1 General

To give an impression of the signal quality and visually exemplify the significance of MAE, examples of data of MAE in the range 0 – 10 μV are given in Fig 1. Each signal was randomly chosen from a randomly chosen EEG, with the exception that drawn examples containing no obvious biological signal (e.g., square waveform signal) were discarded and a new example was drawn.

**Fig. 1.**
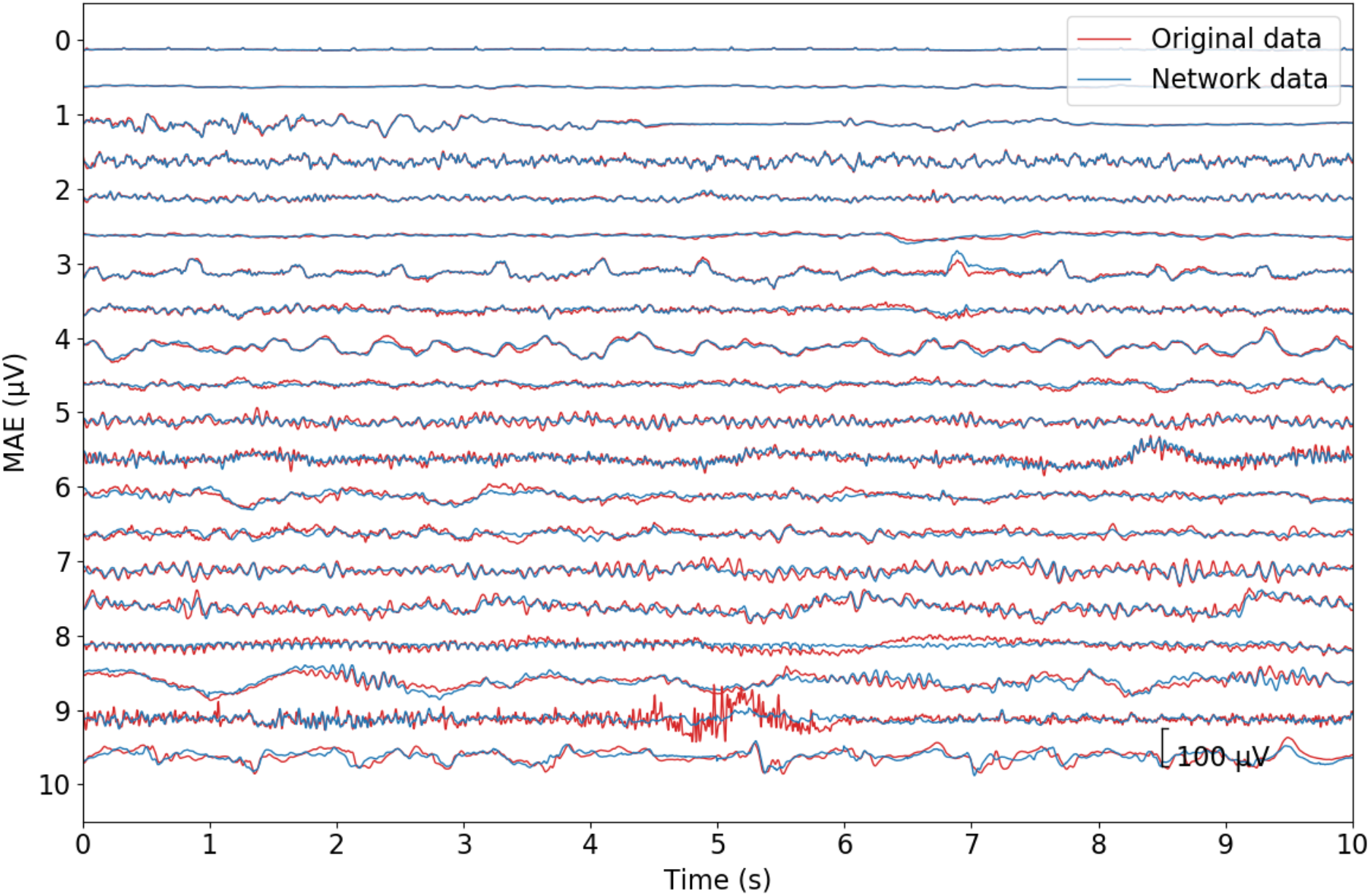
Examples of recreated EEG signals with different MAE to illustrate the visual significance of the measure. Signals were randomly selected with increasing MAE, spanning the interval 0 – 10 μV. The original signal is in red and the signal recreated by GN1 is in blue.

The electrodes relative positions on the scalp surface were approximated by a 2-dimensional grid geometry with their respective median value of the MAE (Fig. 2). For the GN1 network, a gradient was seen with increasing error in central to peripheral direction, which was also seen for the GN3 network, although not as prominent. There was also a gradient with increasing errors in posterior to anterior direction, which was seen in all networks.

**Fig. 2.**
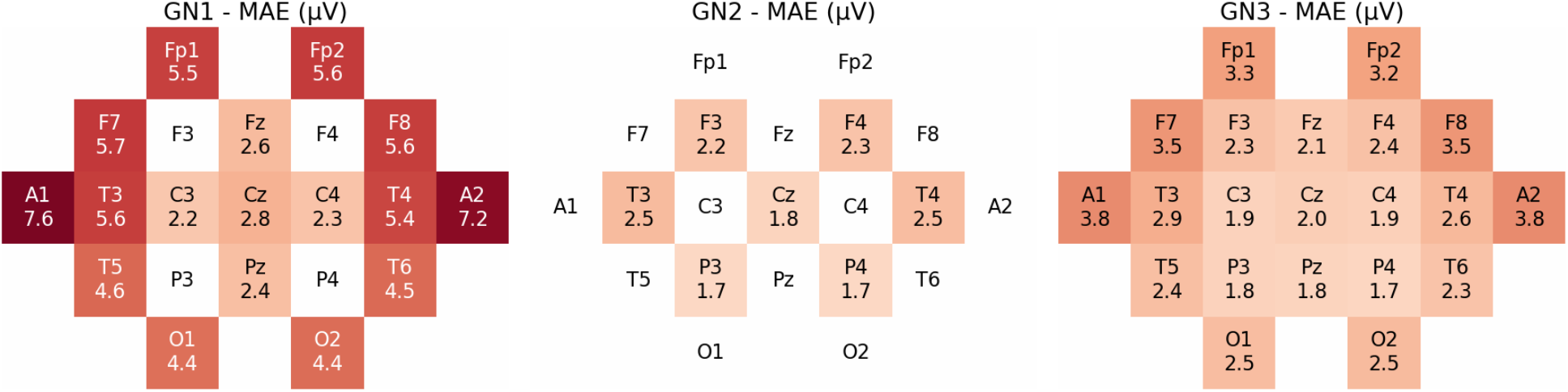
Values of MAE for each electrode for the networks GN1, GN2 and GN3, left to right. The color code from white to dark red represents the MAE values in the range 0 – 8 μV. Each value is based on 12,800,000 data points.

### 3.2 Comparison with spherical spline interpolations

For the GN1 network, the ‘constrained optimization by linear approximation’ algorithm used in the MNE package usually did not converge. This was expected since the problem of interpolating 17 values from 4, where in addition some of the values to be interpolated are spatially distant, is ill posed.

Overall, most measures showed a better performance for the convolutional networks compared to the spherical spline interpolation; 15 out of the 18 calculated values were significantly better (Fig. 3). However, the absolute difference between recreating the signals with convolutional networks or by interpolation were relatively small.

MAE, FD and R were all in favor of the networks. For KL the divergence was smaller for the GN1 and GN2 but larger for the GN3. C and P were better for GN2 and GN3, with higher coherence and less difference in phase compared to interpolation, but the opposite applied to GN1.

**Fig. 3.**
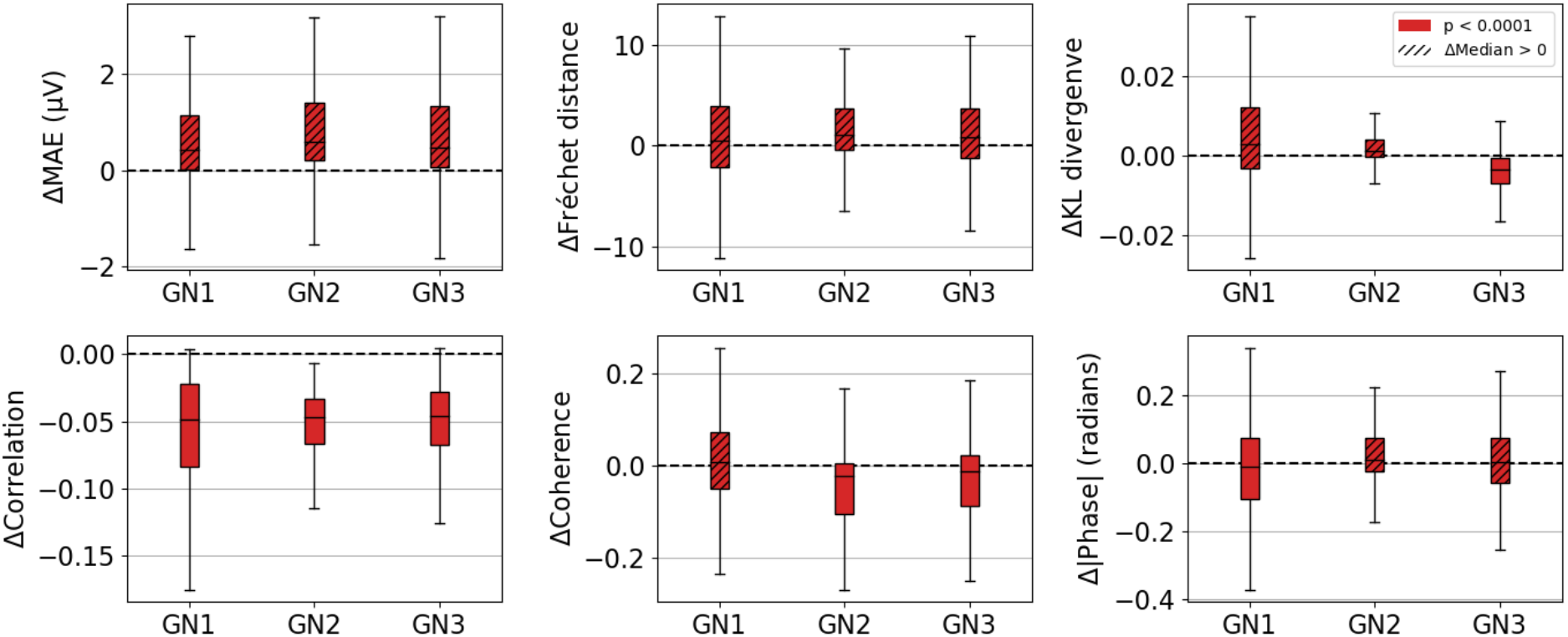
Box-and-whiskers plots (minimum, maximum, median, and first and third quartiles) of the difference of the median between the CNNs and interpolation for all measures. Values from CNNs were subtracted from their paired interpolated values. The number of channels c per network were 17, 7 and 21 for GN1, GN2 and GN3, respectively. The number n of values per channel were 5,000, 500, 100, 100, 5,000×41 and 5,000×41 for MAE, FD, KL, R, C and P, respectively. Each box-and-whisker in the graphs were calculated from c×n values.

Performing predictions using a GPU gave processing times that were shorter than for interpolation (using CPU); Table 5. Predicting with CPU increased the processing times by a factor of 3 – 11, depending on network used.

**Table 5.** Time in seconds (s) for evaluating 5,000 examples with predictions by the networks GN1, GN2 and GN3 or by spherical spline interpolation for the corresponding problems.

|  | GN1 | GN2 | GN3 |
| --- | --- | --- | --- |
| Network, GPU (s) | 41 | 29 | 85 |
| Network, CPU (s) | 476 | 319 | 235 |
| Interpolation (s) | 113 | 41 | 95 |

### 3.3 Visual tests

The randomization gave a distribution of 34, 33 and 33 examples of original, network generated and spline interpolated data with 88, 82 and 52 percent rated as real (Fig. 4), respectively. There was no difference between the real and network generated data, whereas both performed better when compared to interpolated data.

**Fig. 4.**
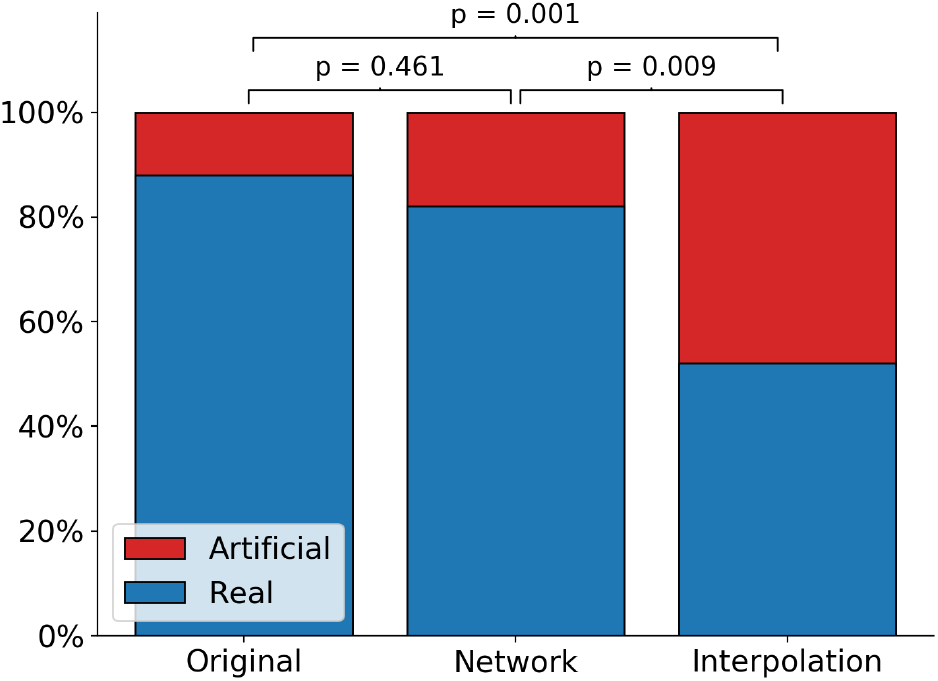
Overall results of the visual assessment. The bars represent, from left to right: original data, data generated by GN1 and spline interpolation. Blue and red indicate the percentage of the examples that raters assessed as real or artificial, respectively. Chi-square statistics were 10.8 (top bracket, p = 0.001), 0.5 (lower left bracket, p = 0.461) and 6.8 (lower right bracket, p = 0.009).

**Fig. 5.**
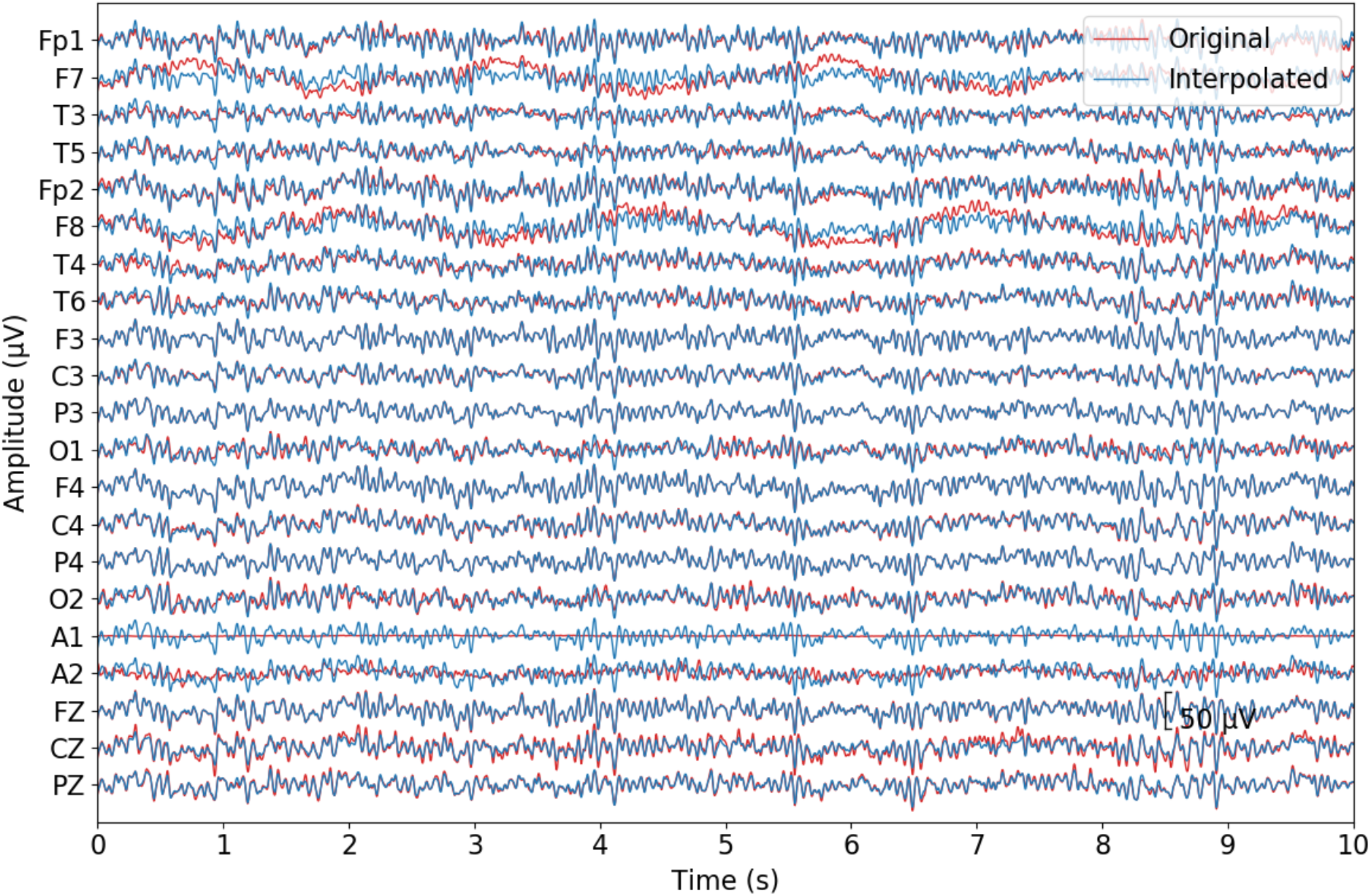
Example of interpolated data (blue) and original signals (red), which all raters assessed as artificial.

For the original data there were none-, for the GN1 network two- and for the interpolation eight examples where all raters were in full agreement as to them being artificial. These examples were visually inspected by author MS (specialist in clinical neurophysiology), and the impression was that they looked cleaner than the originals, i.e., fewer artifacts, and less variation resulting in already monomorphic signals looking even more monomorphic. This was in line with spontaneous comments by some raters after the test, and some also thought that some examples looked ‘too good’.

### 3.4 The effect of the number of subjects

An increase in performance was seen by using more subjects for training. The largest effect was seen for up to 100 subjects, after which only a small additive effect was seen (Fig. 6). However, the magnitude of the improvement created by adding subjects, depended on the network and parameter assessed.

**Fig. 6.**
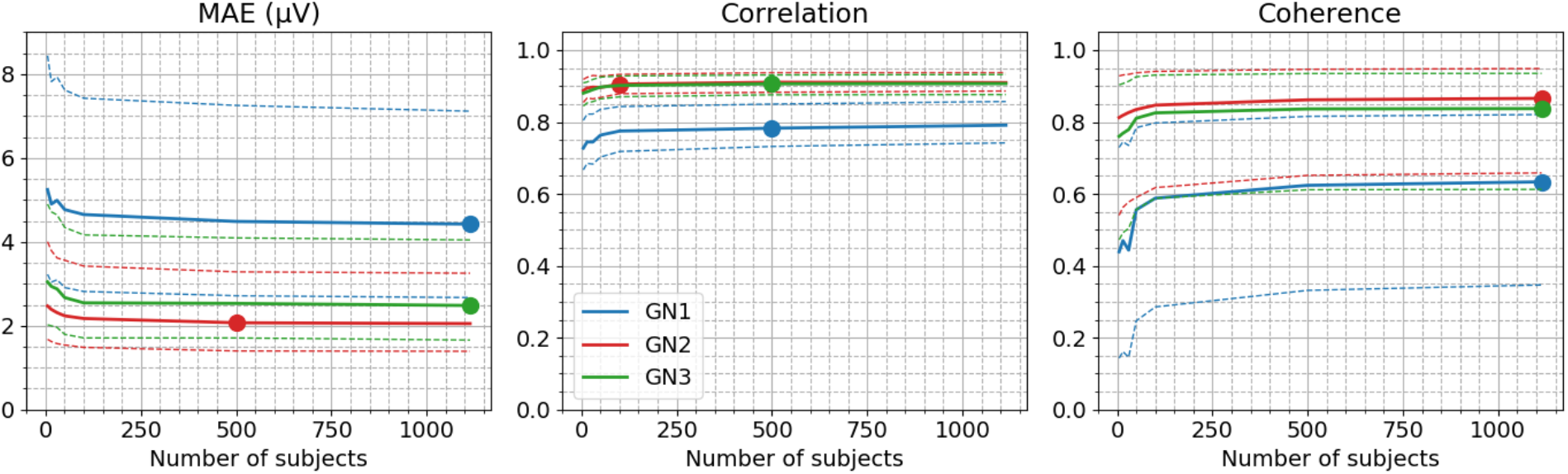
MAE, correlation and coherence as a function of the number of subjects (5, 15, 30, 50, 100, 500, 1,114) used for training of the networks GN1 (blue), GN2 (red) and GN3 (green). Solid lines are the median, dashed lines the 25^th^ and 75^th^ percentiles, and the filled circles mark the points up to which there is a significant effect of adding more subjects. (Kruskal-Wallis H test were used for multiple comparisons and post hoc analyses were performed by Dunn’s multiple comparison test using an adjusted p-value corresponding to 0.05.)

## 4 Discussion

We showed that the convolutional neural networks could recreate EEG signals with a higher precision and produce signals with a more credible visual appearance than one of the most commonly used techniques, spherical spline interpolation. Previous studies (Corley and Huang, 2018; Kwon et al., 2019) have indicated a performance superior to different interpolation methods using smaller data amounts, but comparisons are difficult due to large methodological discrepancies. In the work by Corley et al. (2018), the units of MAE values are not given. Kwon et al. (2019) shows high correlations (up to 0.99) but their evaluation data likely have a better signal-to-noise level since it consisted of either simulated or trial averaged ERP data. Here we used randomly selected examples from a large pool of unsorted real data.

Three different variations of a network architecture were trained. The first network was trained to solve a severely underdetermined problem with the input outnumbered by the output signals, of which some were spatially distant. The specific input electrodes (F3, F4, P3 and P4) used for this network were chosen since they often are used for long term monitoring in intensive care units. As a comparison, the second network was trained under more favorable conditions having an even spatial distribution of input signals, the output signals were all surrounded by known signals and were fewer in numbers. The third network, in addition learned to detect and recreate missing signals dynamically, demonstrating the versatility of deep neural networks and the potential for automating multi-step EEG processing.

In this work, the focus was not on testing a large variety of architectures and parameters. We used a fairly simple and straight forward network architecture without extensive finetuning and still outperformed the standard method. This means that there are probably better network architectures and hyperparameters so that the performance might be further improved.

Whether the processing times are perceived as acceptable depends, apart from the network used, on the available hardware and the amount of data to be interpolated. Using CPU for a standard recording of 20 min consumed 5.6 – 11.4 s for prediction, with the computer used for the tests. That would to many be seen as an acceptable duration.

The amount of data used for our study was relatively large and the networks were trained with an even distribution of examples with respect to the number of subjects. The largest increase in performance was seen in the range 5 – 100 subjects, with only slight increases thereafter. This corresponded to a maximum of 111 – 278 hours of training data, a number that might be reduced given the overlapping training schedule. The impact of the number of subjects vs. the amount of data was not investigated but it is reasonable to assume that the amount of data per subject also influence the results, including the number of subjects needed to saturate the improvement. The distributions of normal vs. pathological and of physiological variations in the data as well as the occurrence of artifacts, were largely unknown and not controlled for in the analysis. If a data set used for training has an unequal distribution in terms of recording length per subject it may be beneficial to prioritize training evenly with respect to the number of subjects. Generally, short recordings tend to yield small-, long recordings large variability. Using data with well described distributions and balanced data sets, may improve performance further. However, assessing a data material of the size used here in such detail, is an extensive undertaking.

Even though the dataset used in this work was large, this does not guarantee that the results generalizes well to other datasets. If the networks trained here are applied to-, or if new networks are trained using data from a different database, the performance and the number of subjects needed to saturate the increase in performance may differ compared to our results.

Evaluating data of different character may reveal pros and cons of the method, e.g., how different types of artifacts affect the results, and how wave transients and spatially restricted activity are reproduced. It is reasonable to assume that signals are more likely to be reproduced correctly if they are synchronized and have a wider spatial distribution, with the opposite being true for localized and random signals, since the prediction of one signal from another depend on the existence of information of the first signal in the second (Lauritzen, 1974). Since synchronization is a hallmark of cortical activity (Nunez and Srinivasan, 2006a), the prediction of the activity in one location from another is plausible. However, since small variations in timing have a higher probability of causing destructive interference for higher frequencies, maintain phase-synchronization over longer distances maybe harder for these frequencies. Moreover, there is evidence that higher frequencies have a more limited spatial synchronization due to resonance phenomenon (Nunez and Srinivasan, 2006b). This could mean that predicting high frequency activity at low electrode densities may be harder. In addition, there are many types of possible artifacts that may be superimposed on the cortical signals and these usually have certain characteristics with respect to morphology, spatial distribution, synchronization and randomness (Tatum et al., 2018). One common source of artifacts is muscle activity. This generates a relatively stochastic electrical signal (Reaz et al., 2006), resulting in an artifact with a random character (an example of a muscle artifacts, clearly not recreated by the network, is seen in Fig.1 at a MAE of 9 μV). Frontal and anterior temporal electrodes have more muscle artifacts. Possibly in line with this, a posterior to anterior gradient was noted, with larger errors in the frontal quadrants (Fig. 2). Eye movements, a low frequency, more organized artifact, naturally affects frontal electrodes the most and likely contributed to the error gradient. Probably adding to this is the higher degree of synchronization over posterior parts due to posterior dominant rhythms. There was also an increasing error in radial direction (Fig. 2). This is probably an effect of distance, due to the larger error when predicting values over longer distances.

In addition to evaluating how the networks recreates various waveforms, e.g., epileptiform activity, the clinical usefulness has to be evaluated. Theoretically a full EEG montage should give a higher sensitivity for the visual detection of many pathological signals compared to a montage with reduced electrode number. Whether the networks recreate the EEG signals faithful enough needs to be evaluated by comparing clinical diagnosing using a full EEG montage recreated by a network, an reduced EEG montage with fewer electrodes and a full EEG montage of original data as baseline.

In this work, we showed that networks can upsample and restore signals. This suggests that the networks are robust to noise or missing information, which have an impact on how they perform during detection or classifications tasks. It is not known if the networks learned the statistical distributions of the electrical fields or if they just learned a simpler way of interpolating the values. The visual tests suggest that the signals recreated by the network have a more natural variation, which may imply that the former is the case. Ordinary interpolation does not add information, so upsampling data by interpolation should not increase the accuracy of neural networks. Adding a general knowledge of the statistical distributions could theoretically boost performance. Thus, training networks having a low number of input channels to also learn the distributions of a larger set of channels might accomplish such a performance boost.

## 5 Conclusions

We demonstrate that restoring or interpolating EEG signals can be performed with neural networks with a precision as good as, or better than, spherical spline interpolation and often with a more credible appearance. Processing times are acceptable for standard EEG exams. It may be worth-while using up to 100 subjects for training a network for this type of task. Minor further gains in performance is possible by adding more subjects but may not be worth the effort if data is hard to acquire. There may be room for improvement by testing other network architectures and using a more content-controlled data. The present networks need further evaluation to assess how they reproduce different important wave phenomena, spatial asymmetries and how different artifacts affects the results. The networks developed here could potentially be used clinically to recreate full montage EEGs from four electrode long term monitoring and recreate bad channels to provide a better visualization, but the usefulness of such an application have to be evaluated clinically.

## Acknowledgement

We would like to thank the Nvidia corporation, who donated the Quadro P6000 and Titan X graphics cards used for training the networks.

